# Unraveling the role of *Slc10a4* in auditory processing and sensory motor gating: implications for neuropsychiatric disorders?

**DOI:** 10.1101/2023.03.14.532543

**Authors:** Barbara Ciralli, Thawann Malfatti, Markus M. Hilscher, Richardson N. Leao, Christopher R. Cederroth, Katarina E. Leao, Klas Kullander

## Abstract

**Background:** Psychiatric disorders, such as schizophrenia, are complex and challenging to study, partly due to the lack of suitable animal models. However, the absence of the *Slc10a4* gene, which codes for a monoaminergic and cholinergic associated vesicular transporter protein, in knockout mice (*Slc10a4* ^-/-^), leads to the accumulation of extracellular dopamine. This makes them a potential animal model for schizophrenia, a disorder known to be associated with altered dopamine signaling in the brain.

**Methods:** The locomotion, auditory sensory filtering and prepulse inhibition (PPI) of *Slc10a4* ^-/-^ mice were quantified and compared to wildtype (WT) littermates. Intrahippocampal electrodes were used to record auditory event-related potentials (aERPs) for quantifying sensory filtering in response to paired-clicks. The channel above aERPs phase reversal was chosen for reliably comparing results between animals, and aERPs amplitude and latency of click responses were quantified. WT and *Slc10a4* ^-/-^ mice were also administered subanesthetic doses of ketamine to provoke psychomimetic behavior.

**Results:** Baseline locomotion during auditory stimulation was similar between *Slc10a4* ^-/-^ mice and WT littermates. In WT animals, normal auditory gating was observed after i.p saline injections, and it was maintained under the influence of 5 mg/kg ketamine, but disrupted by 20 mg/kg ketamine. On the other hand, *Slc10a4* ^-/-^ mice did not show significant differences between N40 S1 and S2 amplitude responses in saline or low dose ketamine treatment. Auditory gating was considered preserved since the second N40 peak was consistently suppressed, but with increased latency. The P80 component showed higher amplitude, with shorter S2 latency under saline and 5 mg/kg ketamine treatment in *Slc10a4* ^-/-^ mice, which was not observed in WT littermates. Prepulse inhibition was also decreased in *Slc10a4* ^-/-^ mice when the longer interstimulus interval of 100 ms was applied, compared to WT littermates.

**Conclusion:** The *Slc10a4* ^-/-^ mice responses indicate that cholinergic and monoaminergic systems participate in the PPI magnitude, in the temporal coding (response latency) of the auditory sensory gating component N40, and in the amplitude of aERPs P80 component. These results suggest that *Slc10a4* ^-/-^ mice can be considered as potential models for neuropsychiatric conditions.

## Introduction

Proteins belonging to the superfamily of solute carriers (SLCs), which includes over 400 distinct proteins that vary widely in their expression and function, are one of the main transporter proteins across cell membranes (Pizzagalli et al, 2020; Lin et al, 2015). While SLCs are well known for transport of neurotransmitters such as dopamine, norepinephrine, GABA and glycine (Kristensen et al, 2011), the function of orphan SLCs expressed in the brain is still unclear.

The orphan solute carrier transporter protein SLC10A4 was first identified in brain, placenta and liver (Splinter et al, 2006) and has been shown to be specifically located at presynaptic vesicles of monoaminergic and cholinergic neurons (Larhammar et al, 2015; Burger et al, 2011; Geyer et al, 2008). We have previously shown that *Slc10a4* ^-/-^ mice have increased sensitivity to cholinergic drugs in epilepsy studies (Zelano et al, 2013), disturbed development of the neuromuscular junction (Patra et al, 2014) and a reduction in striatal serotonin, noradrenaline and dopamine concentrations, with concurrent slower dopamine clearance rate, resulting in the accumulation of extracellular dopamine (Larhammar et al, 2015).

*Slc10a4* ^-/-^ mice have also been shown to be hypersensitive to psychostimulants such as amphetamine and the monoamine oxidase inhibitor tranylcypromine, suggesting that SLC10A4 protein influence storage, release and/or uptake of monoamines by other transporters (Larhammar et al, 2015). Still, there is no evidence that SLC10A4 protein can transport neurotransmitters by itself (Schmidt et al, 2015), and *Slc10a4* ^-/-^ mice have shown reduced dopamine levels in the striatum, and reduced acetylcholine content in the hippocampus and brainstem (Melief et al, 2016). The expression of *Slc10a4* mRNA in the striatum, ventral tegmental area, substantia nigra, locus coeruleus, dorsal raphe and spinal cord motor neurons indicates that this protein may have a regulatory role in several psychiatric conditions since dopaminergic, noradrenergic, serotonergic and cholinergic neurons express SLC10A4 (Larhammar et al, 2015; Geyer et al, 2008).

Psychiatric disorders are difficult to study in animal models, due to the human nature of symptoms that are not easily quantified. However, deficiency in sensorimotor gating and auditory sensory gating, especially auditory filtering of repetitive stimuli, are neurophysiological hallmarks of schizophrenic patients (Brockhaus-Dumke et al, 2008; Javitt and Sweet, 2015; Shen et al, 2020; Freedman et al, 2020) and have shown some reproducibility in animals following drug administration (Mansbach et al, 1988; Miller et al, 1992; Swerdlow, 1994; Winship et al, 2018). Furthermore, several genetic manipulations to model positive symptoms of schizophrenia have been developed, and are often validated by altered locomotor activity and disruption of prepulse inhibition (van den Buuse, 2009; Afshari et al, 2017). Schizophrenia is characterized by dopamine dysregulation, which plays a crucial role in the development of the disease. Studies have shown that the efficacy of antipsychotic medications directly correlates with their affinity to dopamine receptors (Creese et al, 1976; Seeman and Lee, 1975). In addition, glutamate dysregulation is also implicated in schizophrenia positive symptoms. For instance, ketamine, a dissociative anesthetic and a non-competitive antagonist of the N-methyl-D-aspartate (NMDA) receptor, produces schizophrenia-like positive symptoms in healthy humans after a subanesthetic dose (Krystal et al, 1994; Beck et al, 2020) and in mouse models (Connolly et al, 2004; Ehrlichman et al, 2009). However, there is a need for better animal models to study complex neurological conditions such as schizophrenia.

In this study, we aimed to investigate the role of *Slc10a4* in sensorimotor gating, and in auditory sensory gating and its potential interaction with ketamine. We measured auditory event-related potentials in the dorsal hippocampus in response to paired clicks in *Slc10a4* ^-/-^ mice and wildtype littermates. Specifically, we hypothesized that the lack of *Slc10a4* would impair auditory sensory gating and that *Slc10a4* ^-/-^ mice would be more sensitive to ketamine’s impairment of auditory gating.

## Methods

### Animals

Mice were housed as approved by the animal care unit of Uppsala University and experiments were conducted according to Swedish guidelines and regulations, and European Union legislation (ethical permits C248/11 and C3/12).

Male and female mice (12-24 weeks old) from heterozygote breedings of *Slc10a4* ^+/-^ mice of 129/SvEvBrd background (Texas A&M Institute for Genomic Medicine, TX, USA) were inbred with C57BL/6 for at least three generations to obtain null mutant *Slc10a4* ^-/-^ and wild-type (WT) littermates on a stable genetic background. Ten *Slc10a4* ^-/-^ mice and 6 WT littermates were used in this study. One WT and one *Slc10a4* ^-/-^ mouse were excluded due to the poor electrophysiological signal after electrode implantation, and 3 *Slc10a4* ^-/-^ did not survive the implantation surgery, leading to a total of 11 mice (6 *Slc10a4* ^-/-^ mice and 5 WT) reported in aERP experimental procedures. For prepulse inhibition experiments a separate set of animals were used (8 WT and 8 *Slc10a4* ^-/-^ mice).

### Hippocampal chronic electrodes implant

Mice were anesthetized with inhalation of isoflurane at an initial dose of 4 %, then 1-2 % during the remainder of the procedure. The *Slc10a4* ^-/-^ animals were more sensitive to anesthesia and 3 mice died during/after surgery. Lowering the dose to 1 % isoflurane proved to be adequate anesthesia of *Slc10a4* ^-/-^ mice.

For intrahippocampal electrode implantation animals were placed on a heating pad and immobilized in a standard stereotaxic frame. Eye gel (visgel) was applied to avoid corneas from drying. Iodine and lidocaine was applied, the skin opened and four holes were drilled into the skull with a dental drill. One hole for the chronic 16 channel electrode (CM16LP, silicon-substrate multichannel A1×16-electrode probe, weight 160mg, recording sites spaced 100 *μ*m apart distributed along a single shank, NeuroNexus, Ann Arbor, MI, US) at coordinates relative to bregma; AP: −2.18 mm, ML: 2.0 mm and DV: 2.5 mm, corresponding to the dorsal left hippocampus. Three micro-screws were placed into the remaining holes (two were pre-soldered to the electrode serving as reference and ground) securing the cap of dental cement to the skull that fixates the chronic electrode. The ground and reference electrodes were placed in the occipital bone or contralateral temporal bone, with some variability due to the length of silver wire soldered to the micro screw. At the end of the surgery mice were rehydrated with a subcutaneous injection of saline, maintained on a heating pad and monitored until fully awake. Mice were allowed to recover for 2 weeks before recordings.

### Auditory event-related potentials

Animals were assessed for deficits in sensory filtering using a paired-click paradigm for recording auditory evoked potentials (Siegel et al 2005; Bickel et al 2007; Ciralli et al 2022, *pre-print*). Mice were placed in a standard plastic cage bottom inside a custom-built sound shielded chamber fitted with LED lights and a digital camera to record behavior. The implanted electrode was attached to an Intan chip connected to an Intan 16 channel amplifier based on the RHA2000-EVAL board (Intan technologies, Los Angeles, CA, US). Animals were left 15-30 min to acclimatize to the environment and head connection before sound presentation started. A loudspeaker (Tweeter, R2904/700009, Scan-Speak, Denmark) was placed above the cage and driven by an amplifier (PPA200, Pyle Audio Inc., Brooklyn, NY, US). Sound stimulation, as paired clicks (80 dB sound pressure level, 10 ms duration, 500 ms inter-click interval, randomized at 2-13 s, 70 repetitions), was generated in Matlab (Mathworks). A 68-Pin data acquisition device (SCB-68A, National instruments) was used to output TTL signals indicating the timing of the randomized paired-click stimuli, which were then recorded in synchrony with the electrophysiological signal with the same Intan RHA2000-EVAL board. An impedance test was routinely recorded to confirm appropriate electrode resistance at the 16 channels.

### Prepulse Inhibition (PPI) and Acoustic Startle Reflex

8 *Slc10a4* ^-/-^ and 8 WT mice were examined using Prepulse Inhibition (PPI) testing. Four sessions with four mice each were run. Startle, startle habituation and PPI of startle were measured using four units of an automated system (SR-LAB, San Diego Instruments, USA). The mice were placed in clear, evenly perforated Plexiglass cylinders, closed on each end and acoustic stimuli were delivered over 68 dB background noise through a speaker in the ceiling of each box. The test sessions consisted of 96 trials with an inter-trial interval between 10 and 25 s. The first and last 16 trials consisted of single 20 ms 105 dB pulse alone startle stimuli. The middle 64 trials consisted of random delivery of 16 105 dB startle pulse-only stimuli, 16 trials of pre-pulse only (75 dB) and 32 prepulse trials. The total of 48 105 dB startle pulse trials was expressed as three blocks of 16 and used to determine startle habituation. Prepulse trials consisted of a single 105 dB pulse preceded by either a 30 ms or a 100 ms interstimulus interval (ISI) by a 4 ms non-startling stimulus of 5 or 15 dB over the 70 dB baseline. Startle responses were converted into quantitative values by a piezo-electric accelerometer unit attached beneath the platform.

### Pharmacology

Auditory event-related potentials were recorded under three conditions; intra peritoneal (i.p.) injection of saline (100 *μ*l, internal control), low dose ketamine (5 mg/kg, 100 *μ*l) and subanesthetic dose of ketamine (20 mg/kg, 100 *μ*l).

### Histology

The electrode tract was routinely confirmed post hoc. After all experiments, mice were transcardially perfused with PBS 1 % followed by PFA 4 %. The brain was dissected and sliced into sections of 100-150 *μ*m thickness using a free-floating vibratome. The tissue was then stained with 4’,6-diamidino-2-phenylindole (DAPI) for visualization of the hippocampal layers.

### Data analysis

All mice were video-recorded from the top for one minute before each ERP recording, and mouse position in the video was tracked after the experiments were done. The instantaneous distance was calculated as the euclidean distance between consecutive frames; and the average distance was calculated as the mean of the instantaneous distance.

Auditory ERP analysis was performed similarly to previously described (Ciralli et al, 2022, *pre-print*). Before extracting auditory event-related potentials from the extracellular recordings, broken channels were identified for each animal and removed from further analysis. Auditory event-related potentials were extracted by filtering the recorded signal with a 3-60 Hz 2nd-order bandpass butterworth filter; then slicing 0.5 s before and 1 s after the first click onset; and averaging the resulting 70 slices (one for each pair of click). Next, in order to consistently compare responses from the same approximate hippocampal layer (Scheffer-Teixeira and Tort, 2017), the phase-reversal was identified as the pair of adjacent channels with the largest phase difference, and the channel within this pair with the most negative peak is selected for further analysis. The N40 was considered as the minimum peak within 10-60 ms after the click stimulus, while P80 was considered as the maximum peak within 20-100 ms after the sound click. The components amplitudes were determined as the peak voltage subtracted by the baseline, which is the average of 200 ms preceding the sound onset. The components latencies were determined as the time where the peak was detected after the sound click onset. The percentage of second click suppression and delay (for amplitude and latency measurements, respectively) were calculated as

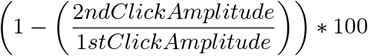

with error whiskers representing the standard error of the mean (s.e.m) for all figures.

The effect of genotype and treatment (for total distance, average distance, ERP ratios) or genotype, treatment and click repetition (for ERP amplitudes and latencies) were evaluated using mixed models ANOVA, and pairwise comparisons were evaluated with t-test, corrected for unequal variances when needed (evaluated with the Levene test). Where data failed to comply with normality (evaluated with the Shapiro-Wilk test), effects were evaluated using Friedman test (for treatment and click) or Kruskal-Wallis test (for genotype), and pairwise comparisons were evaluated using Wilcoxon signed-rank (between treatments or clicks) or Mann-Whitney U test (between genotypes). All multiple comparisons had their p-value corrected using Holm’s correction.

PPI analysis was performed in Matlab 2022b after exporting startle responses from SR-LAB. Percentage prepulse inhibition (%PPI) was calculated as

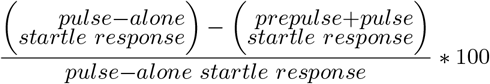

and effect of genotype, interstimulus interval and stimulus intensity were evaluated using mixed-models ANOVA. Pairwise comparisons between stimulus interstimulus intervals and stimulus intensities were evaluated using paired t-test, while pairwise comparisons between genotypes were evaluated using Tukey’s HSD.

### Software availability

Mouse positions in videos were tracked using the Mice Profiler Tracker plugin (de Chaumont et al, 2012), and all electrophysiological data was recorded using the RHA2000-Eval System. All data analysis was performed using SciScripts (Malfatti, 2023), Scipy (Virtanen et al, 2020) and Numpy (Harris et al, 2020). All plots were done using Matplotlib (Hunter, 2007); schematics were done using Inkscape (Inkscape Project, 2022) and merged into the plots using svgutils (Geier, 2021). All scripts are available online (Malfatti and Ciralli, 2023).

## Results

To investigate behavioral and sensory gating capability in *Slc10a4* ^-/-^ mice we first evaluated whether their locomotion was altered compared to wildtype littermates (WT). Animals were recorded during 10 minutes after intraperitoneal injection of saline, followed by 5 mg/kg ketamine and 20 mg/kg ketamine (Figure 1 A-B). Individual locomotion was tracked for WT and *Slc10a4* ^-/-^ animals (Figure 1 C-D, respectively). We found an overall effect of ketamine in the mean distance traveled (ANOVA, F(2,22) = 5.191, p = 1.4e-02), indicating that ketamine caused hyperlocomotion independently of genotype. Although wildtype mice showed higher mean values in average distance after being administered 20 mg/kg of ketamine compared to saline (NaCl: 0.110 ± 0.032 cm and Ketamine 20 mg/kg: 0.237 ± 0.061 cm; Figure 1E), no significant difference between treatments was found (t-test eff. Size < 1.058; p > 0.240). Locomotion tracked for *Slc10a4* ^-/-^ animals showed mean changes in similar direction to WT littermates, where ketamine 20 mg/kg showed highest values for locomotion (*Slc10a4* ^-/-^ saline average distance: 0.142 ± 0.033 cm; ketamine 20 mg/kg: 0.178 ±0.071 cm; Figure 1E), however, no significant differences between genotypes or treatments were found (t-test eff. size < 0.863; p > 0.626).

**Figure 1:**
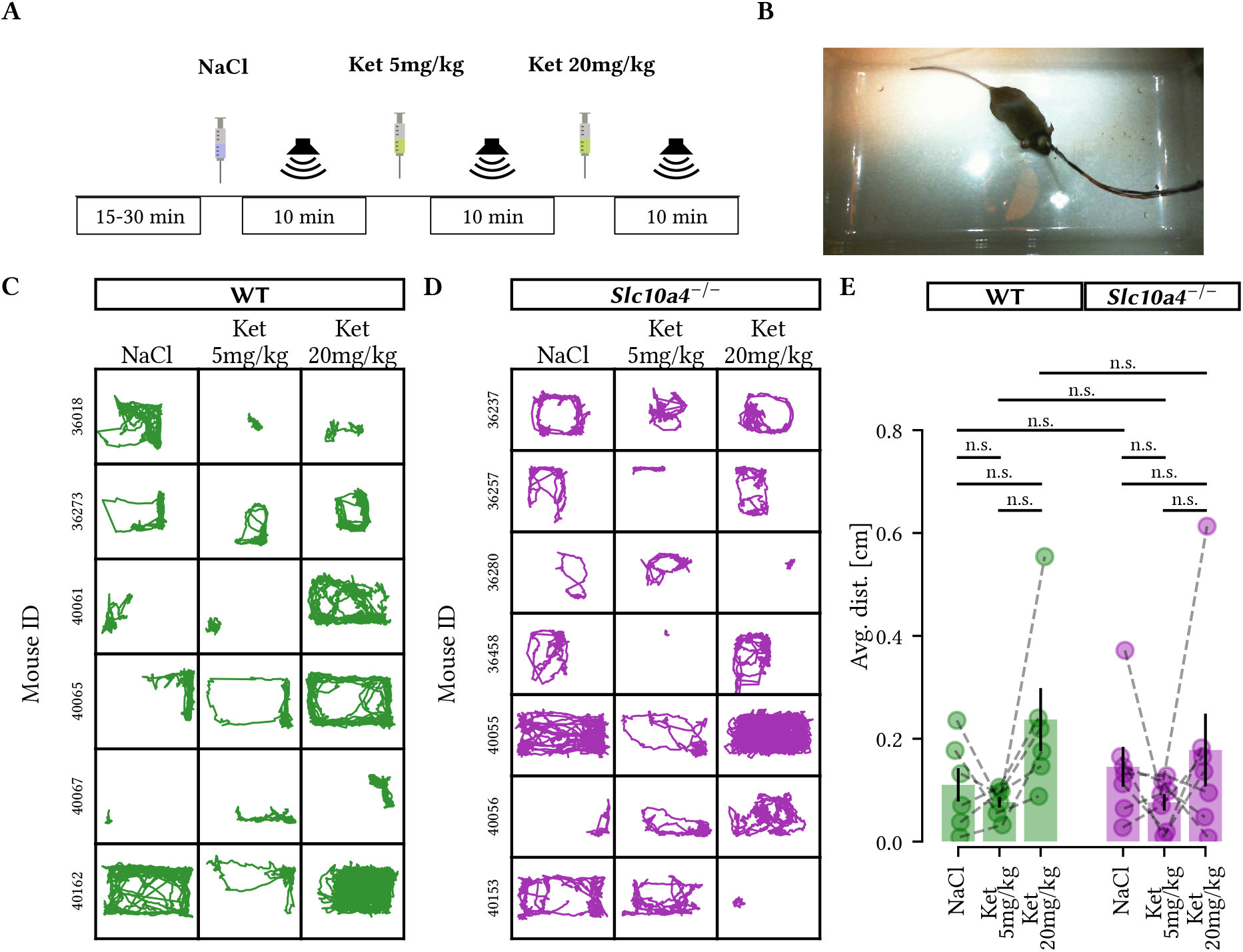
*Slc10a4*^-/-^ do not display affected locomotion compared to control littermates. A) Timeline of the experiments. Locomotion was recorded during 10 minutes after each pharmacological treatment. B) Image of an animal inside the experimental setup. C-D) Individual tracking of locomotor activity in saline, ketamine 5 mg/kg and ketamine 20 mg/kg for WT (green, C) and *Slc10a4* ^-/-^ mice (purple, D). Numbers on the left show the subject ID. E) Average distance traveled show no significant differences between groups (t-test, p > 0.551 for all comparisons) or treatments (t-test p > 0.240 for all comparisons, n = 6 WT and 7 *Slc10a4* ^-/-^).

To robustly compare auditory event-related potentials (aERPs) between different animals and after multiple treatments, animals were implanted in the CA1 region of the left dorsal hippocampus (Figure 2A) and the phase-reversal of aERPs was identified for each recording (Figure 2B). We selected the channel above phase-reversal for analysis of amplitude and latency generating a negative peak at approximately 40 ms (N40) and a positive peak approximately at 80 ms latency (P80) (Figure 2C). Two main parameters were examined: the responses to paired sound clicks (both amplitude in *μ*V and latency in ms, as a readout of sound processing in the limbic system) and the ratio between the second and first click responses, indicating sensory gating. When averaging all aERP waveforms (70 repetitions of paired-clicks per animal for generating a grand average per mouse) from WT and *Slc10a4* ^-/-^ mice, recordings showed that the second click consistently produced a smaller N40 aERP, indicating auditory gating (Figure 2D). Still, visual inspections of waveforms indicated differences across genotypes requiring further dissection.

**Figure 2:**
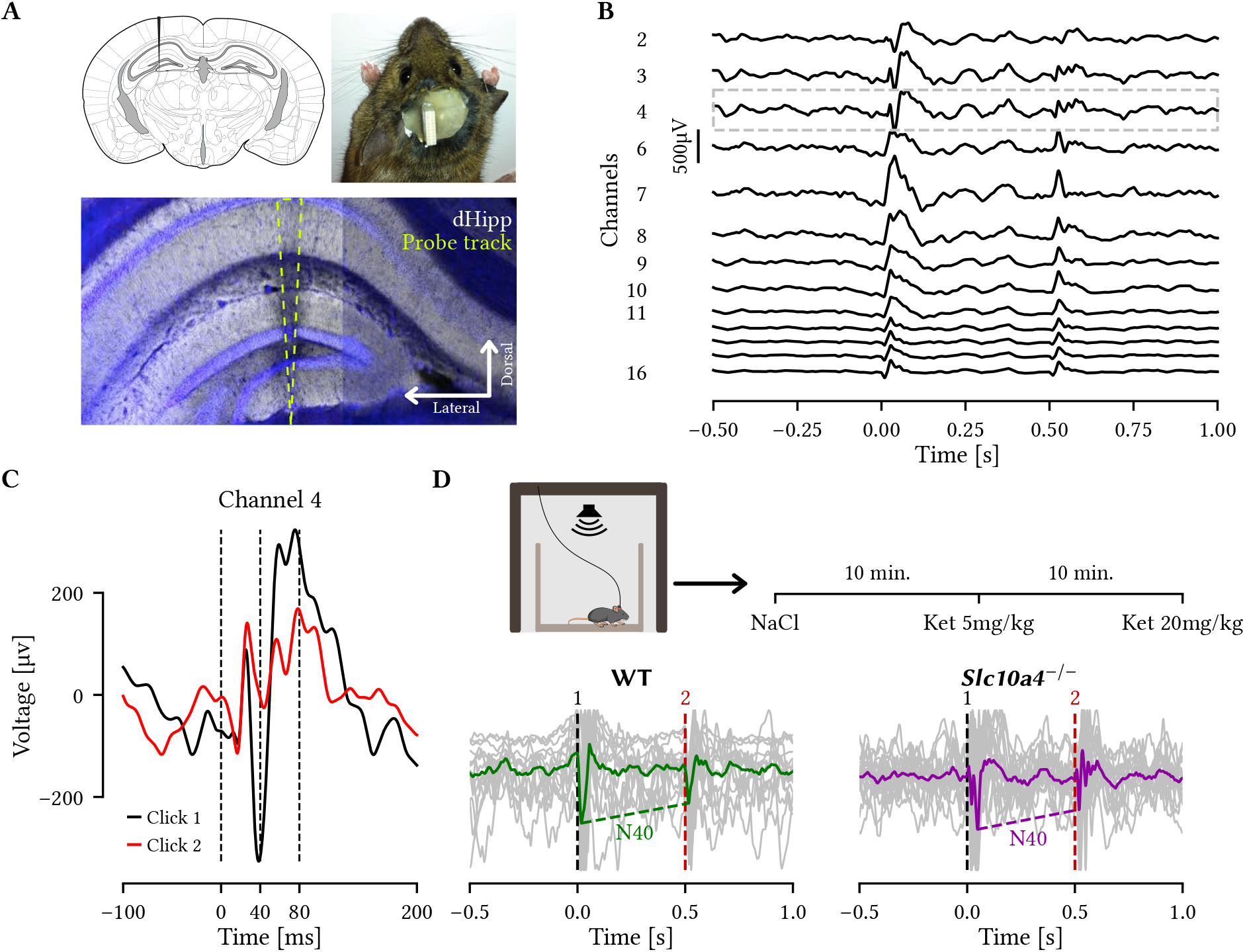
Auditory event-related potentials in WT and *Slc10a4*^-/-^ mice. A) Schematics of the location of silicon probes used for LFP recordings (Top, left; Adapted from Paxinos & Franklin; 2007). Picture of a mouse implanted with a chronic electrode (Top, right). Image of the dorsal hippocampus showing the electrode tract (yellow dashed line) across hippocampal layers (Bottom). B) Representative average aERPs in response to paired clicks from 16 channels at different depths. The channel above phase reversal (gray dotted box) was always used for aERP quantification. C) The reversal channel from ‘B’ at a greater magnification with click 1 (black) and 2 (red) sound responses overlayed. Dashed lines indicating positive and negative peaks at different characteristic latencies (N40 and P80 components). D) Top, simplified experimental timeline. Bottom, average traces of click responses in saline condition for WT (green, n = 5) and *Slc10a4* ^-/-^ animals (purple, n = 6). Superimposed gray traces are the average response of 70 trials from each individual animal, dashed lines indicate the sound stimuli onset and amplitude difference of N40 peaks.

To carry out a detailed quantification of the aERP we first quantified the amplitudes and latency for each N40 response under saline, low-dose ketamine (5 mg/kg) and subanesthetic-dose ketamine (20 mg/kg; Figure 3A). The N40 amplitude of the second click response was as expected significantly decreased compared to the first click in WT animals in the saline condition (t-test eff. size = 1.341; p = 4.5e-02) and also after 5 mg/kg ketamine (t-test eff. size = 0.730; p = 6.3e-03; Figure 3B), but not after administration of 20 mg/kg ketamine (t-test eff. size = 0.155; p = 0.741). Subanesthetic doses of ketamine thereby decreased sensory processing in WT littermates, in agreement with studies using acute ketamine administration as an animal model for psychosis-like reponses (Connolly et al, 2004; Chatterjee et al, 2011). Quantifying N40 components in *Slc10a4* ^-/-^ mice showed no significant difference between first and second click amplitudes in saline (t-test eff. size = 0.511; p = 0.091) or 5 mg/kg ketamine (t-test eff. size = 0.345; p = 0.123), and a trend of second click decrease after 20 mg/kg ketamine (t-test eff. size = 0.356; p = 0.058; Figure 3C). Next, comparing latency of the N40 component showed no differences of the first and second click latency for WT littermates (Wilcoxon signed-rank eff. size < 0.58, p > 0.625; Figure 3D), while *Slc10a4* ^-/-^ mice presented a significant latency decrease in the second click compared to the first click in the saline condition (Wilcoxon signed-rank eff. size = 0.87; p = 3.1e-02; Figure 3E). Despite the low number of animals, all responses were within 2 standard deviations indicating no outliers. Next we analyzed the N40 amplitude ratio, shown as percentage of second peak suppression, and found no significant difference between genotypes or treatments (t-test eff. size < 0.384; p > 0.117; Figure 3F), nor overall genotype effect on amplitude of the N40 response (ANOVA F = 0.012; p = 0.914; Figure 3G). Together, this data indicates that *Slc10a4* ^-/-^ mice show, if any, a mild impairment of N40 suppression in saline condition. In contrast, *Slc10a4* ^-/-^ mice show a significantly shorter latency of the second click response in saline, and a trend in the same direction after 5 mg/kg ketamine.

**Figure 3:**
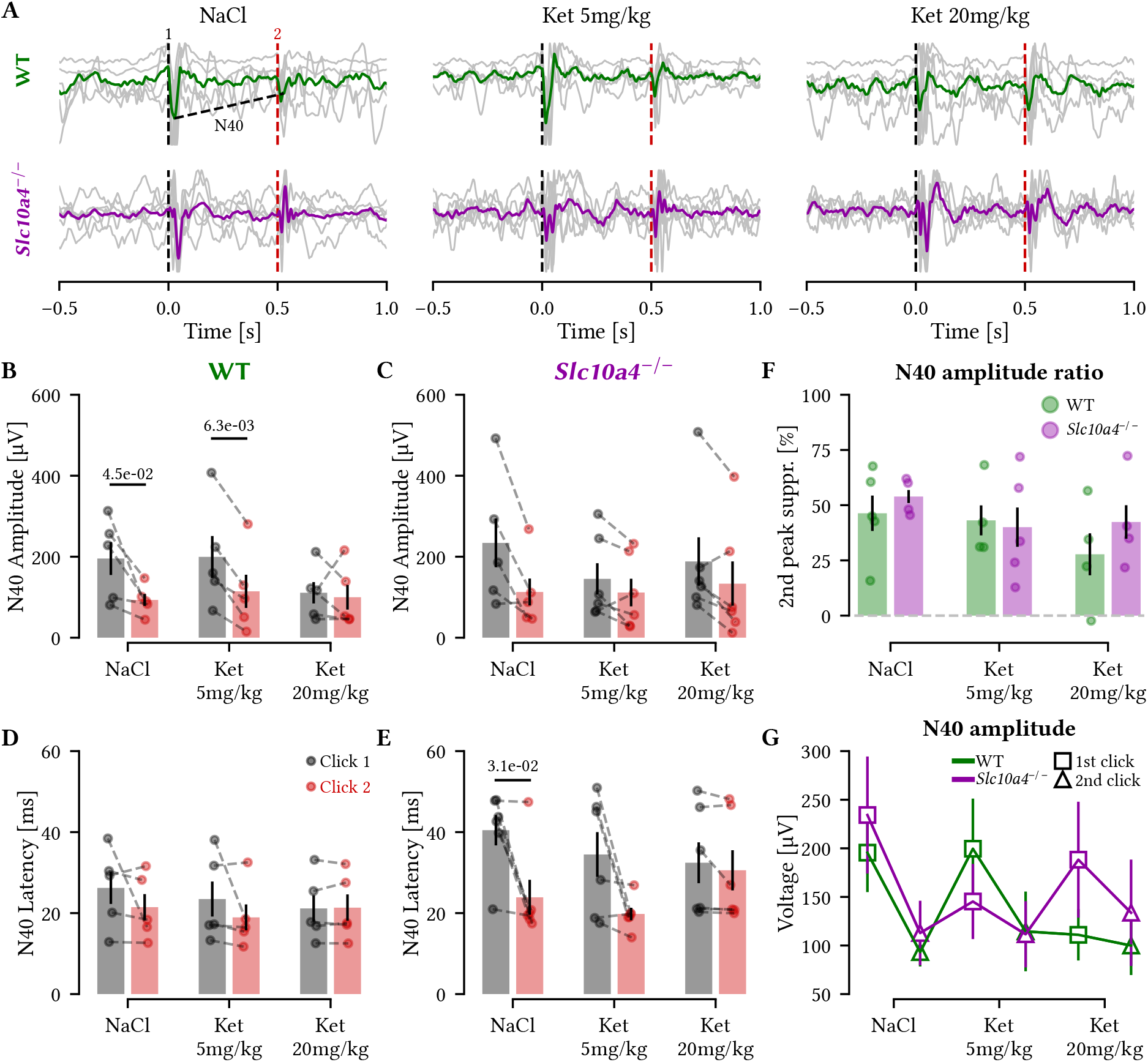
The N40 component is more variable in *Slc10a4*^-/-^ compared to WT animals after saline and ketamine administration. A) Auditory ERP recorded in awake mice in response to saline, ketamine 5 mg/kg and ketamine 20 mg/kg show characteristic suppression of the N40 second click in both WT (top) and *Slc10a4* ^-/-^ (bottom) animals. Gray traces show the average aERP per animal while the green and purple traces show the group average for each treatment. B-C) Quantification of the N40 amplitude in response to the first (gray) and second (red) clicks in different pharmacological treatments for WT (B) and *Slc10a4* ^-/-^ (C) mice. WT mice displayed a decrease in amplitude of the second click in the presence of saline (p = 4.5 e-02, t-test) and Ketamine 5 mg/kg (p = 6.3 e-03, t-test). D-E) WT mice had no significant changes in latencies between clicks (D), while *Slc10a4* ^-/-^ mice presented a significant decrease to the latency of second click in saline condition (p = 3.1 e-02, Wilcoxon signed-rank test; E). F) Percentage of suppression of the second click of the N40 component for WT (green) and *Slc10a4* ^-/-^ (purple) mice. G) Average of the N40 amplitude of WT (n = 5, green) and *Slc10a4* ^-/-^ (n = 6, purple) for each click at each treatment.

Thereafter we investigated whether the positive P80 component of the aERP waveform was affected by the *Slc10a4* ^-/-^ condition or ketamine treatments (Figure 4A). Here, WT littermate mice showed no difference between the first and the second click amplitude after saline, 5 or 20 mg/kg ketamine (t-test eff. size < 0.669; p > 0.148; Figure 4B) even though the second response showed smaller mean values than the first. On the contrary, *Slc10a4* ^-/-^ mice presented a trend of the second click response to be increased compared to the first in saline (t-test eff. size = 0.290; p = 0.063) (Figure 4C). Furthermore, no latency difference was found for WT animals in saline, 5 or 20 mg/kg ketamine between paired clicks (t-test eff. size < 0.531; p > 0.079; Figure 4D). Instead, *Slc10a4* ^-/-^ mice presented a significant decrease of the second click latency in saline condition (t-test eff. size = 1.079; p = 1.6e-02) and 5 mg/kg ketamine (t-test eff. size = 1.877; p = 3.1e-02) but not after administration of 20 mg/kg ketamine (t-test eff. size = 0.375; p = 0.097; (Figure 4E), similar to, and possibly an effect of, the shorter latency N40 component. Although P80 amplitude ratio showed smaller mean values for *Slc10a4* ^-/-^ animals under saline and 5 mg/kg ketamine compared to WT, with the percentage of second peak suppression showing a negative value, indicating second peak enhancement, no significant differences were found between groups (Mann-Whitney-U eff. size < 0.407; p > 0.177) or treatments (Wilcoxon signed-rank eff. size < 0.348; p > 0.438; Figure 4F). However, we found a significant effect of genotype in the P80 amplitude (ANOVA F = 3.798; p = 3.0e-02), with *Slc10a4* ^-/-^ mice displaying consistently increased amplitudes compared to WT littermates, suggesting that *Slc10a4* ^-/-^ animals present abnormal P80 sensory processing (Figure 4G).

**Figure 4:**
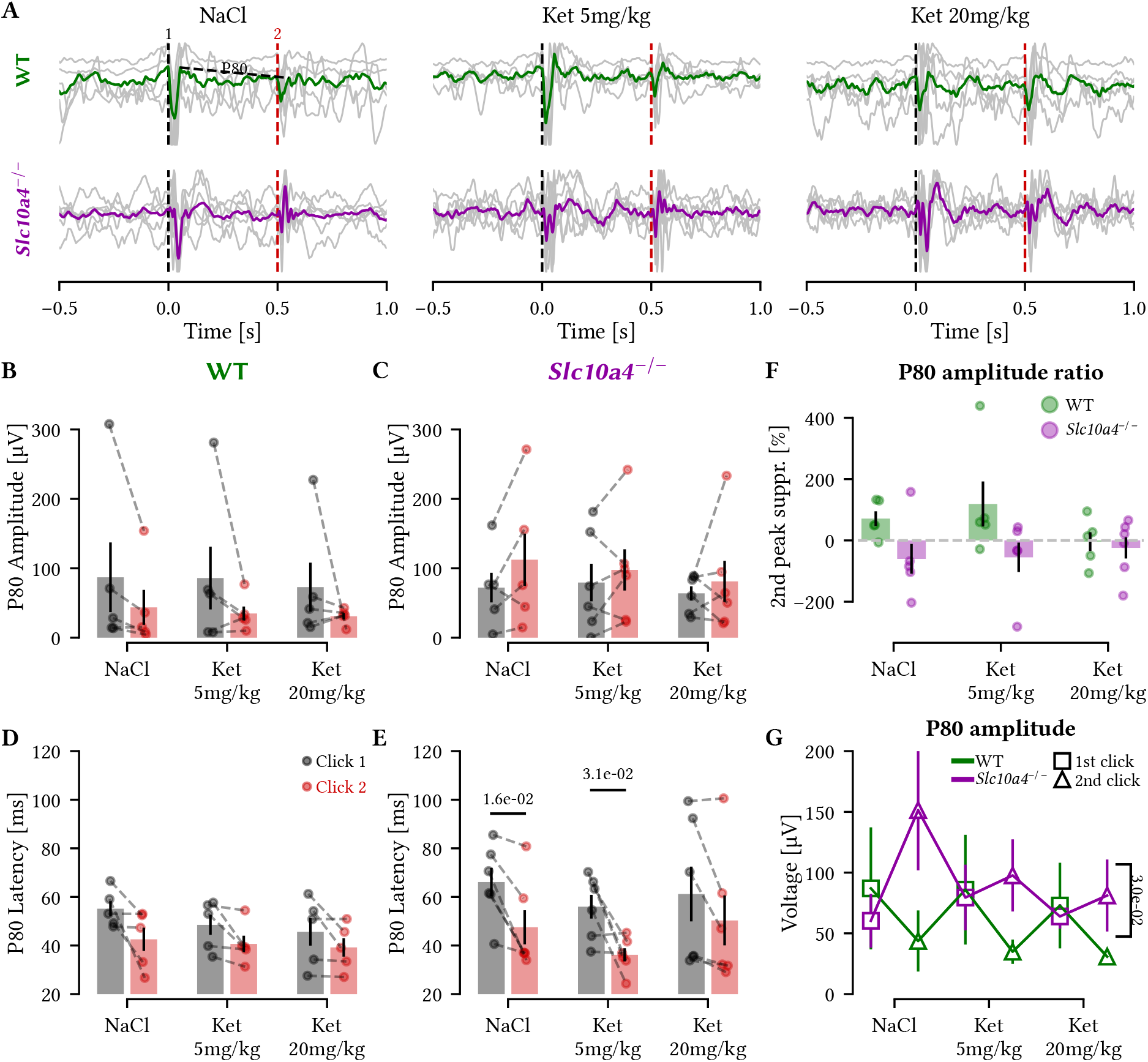
The P80 aERP overall amplitude is increased in *Slc10a4*^-/-^ mice. A) Representative traces (same as Figure 3A) now highlighting the P80 component (vertical black and red dashed lines for first and second click onsets, respectively). B-C) Quantification of the P80 amplitude in response to the first (gray) and second (red) clicks in different pharmacological treatments for WT (B) and *Slc10a4* ^-/-^ (C) mice. The second click showed a trend to increase in *Slc10a4* ^-/-^ mice compared to the first click (t-test eff. size = 1.064; p = 0.063) in NaCl condition. D) The latency of the first and second click was not affected by different treatments in WT mice. E) *Slc10a4* ^-/-^ mice presented a significant decrease to the second click latency in saline condition (p = 1.6e-02, t-test) and ketamine 5mg/kg (p = 3.1e-02, t-test). F) The percentage of P80 second peak amplitude suppression for WT and *Slc10a4* ^-/-^ mice (dashed gray line indicates 0% suppression). G) Group average of the P80 amplitude of WT and *Slc10a4* ^-/-^ for each click at each treatment showing that P80 amplitude in consistently increased for *Slc10a4* ^-/-^ compared to WT animals (p = 3.0e-02, Kruskal-Wallis test). n = 5 WT and 6 *Slc10a4* ^-/-^ mice.

Next we examined the N40 and P80 amplitude responses to each click separately across treatments. For the N40 peak we only found a trend of decrease in click 1 response after 20 mg/kg ketamine in WT mice compared to saline treatment (t-test eff. size = 0.987; p = 0.053), and thereby no significant differences in either first or second click amplitude in both WT and *Slc10a4* ^-/-^ between treatments (Figure 5A). Examining the P80 amplitude separately for the first and second click showed no difference between treatments for neither WT (Figure 5B, top) or *Slc10a4* ^-/-^ mice (Figure 5B, bottom).

**Figure 5:**
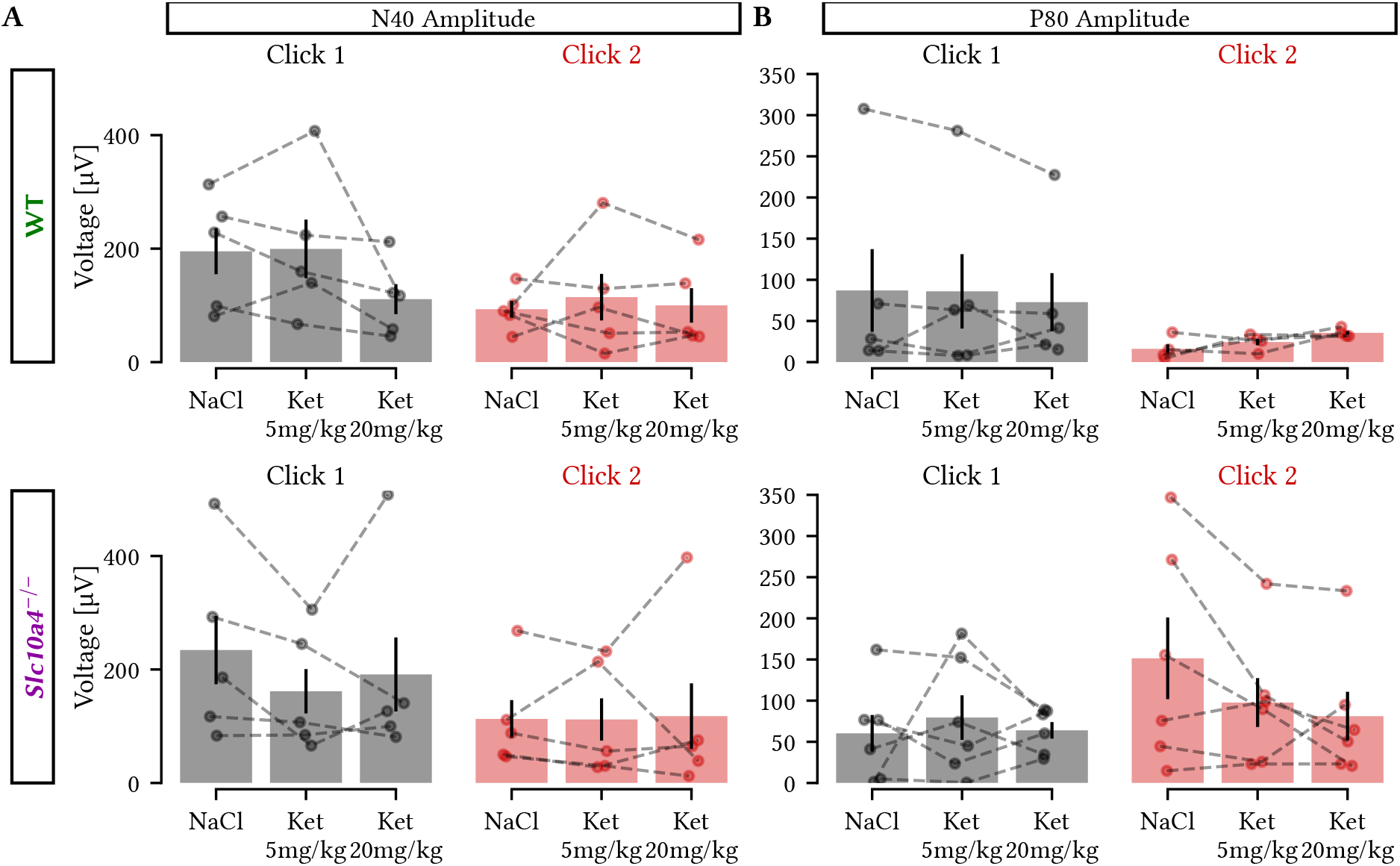
Ketamine does not alter the paired-click N40 and P80 amplitudes compared to saline. A) Comparison of the N40 amplitude in response to the first click (left) and second click (right) after saline, 5 mg/kg ketamine and 20 mg/kg ketamine administration for WT (top) and *Slc10a4* ^-/-^ (bottom) mice. B) Same as ‘A’, but for P80 amplitude. No effect was found across treatments on N40 or P80 amplitude (Wilcoxon signed-rank test). n = 5 WT and 6 *Slc10a4* ^-/-^ mice.

Lastly, focusing on the temporal processing of aERP in WT and *Slc10a4* ^-/-^ mice we analyzed the latency ratio for both the N40 and P80 components. *Slc10a4* ^-/-^ mice animals showed no difference in latency ratio of the two N40 click responses compared to WT littermates in neither treatment (Mann-Whitney-U eff. size < 0.294; p > 0.329; Figure 6A), however, examining the absolute value of the latency in ms, we found that the genotype had a significant effect on N40 latency (Kruskal-Wallis eff. size = 0.142; p = 1.5e-03), where *Slc10a4* ^-/-^ mice showed longer latency, suggesting slower temporal processing in knockout mice (Figure 6B). Analyzing the P80 latency ratio showed no pairwise differences between groups or treatments (t-test eff. size < 0.362; p > 0.237; Figure 6C), and no effect of genotype on P80 latency (ANOVA F = 2.278; p = 0.165; Figure 6D). This is interesting since it shows that the increased N40 latency in *Slc10a4* ^-/-^ mice is not due to a shift in aERP waveform, as this would also alter the P80 component latency, which was not seen. Instead, *Slc10a4* ^-/-^ mice appear to have distinct alterations to aERP latency of the early component of auditory gating.

**Figure 6:**
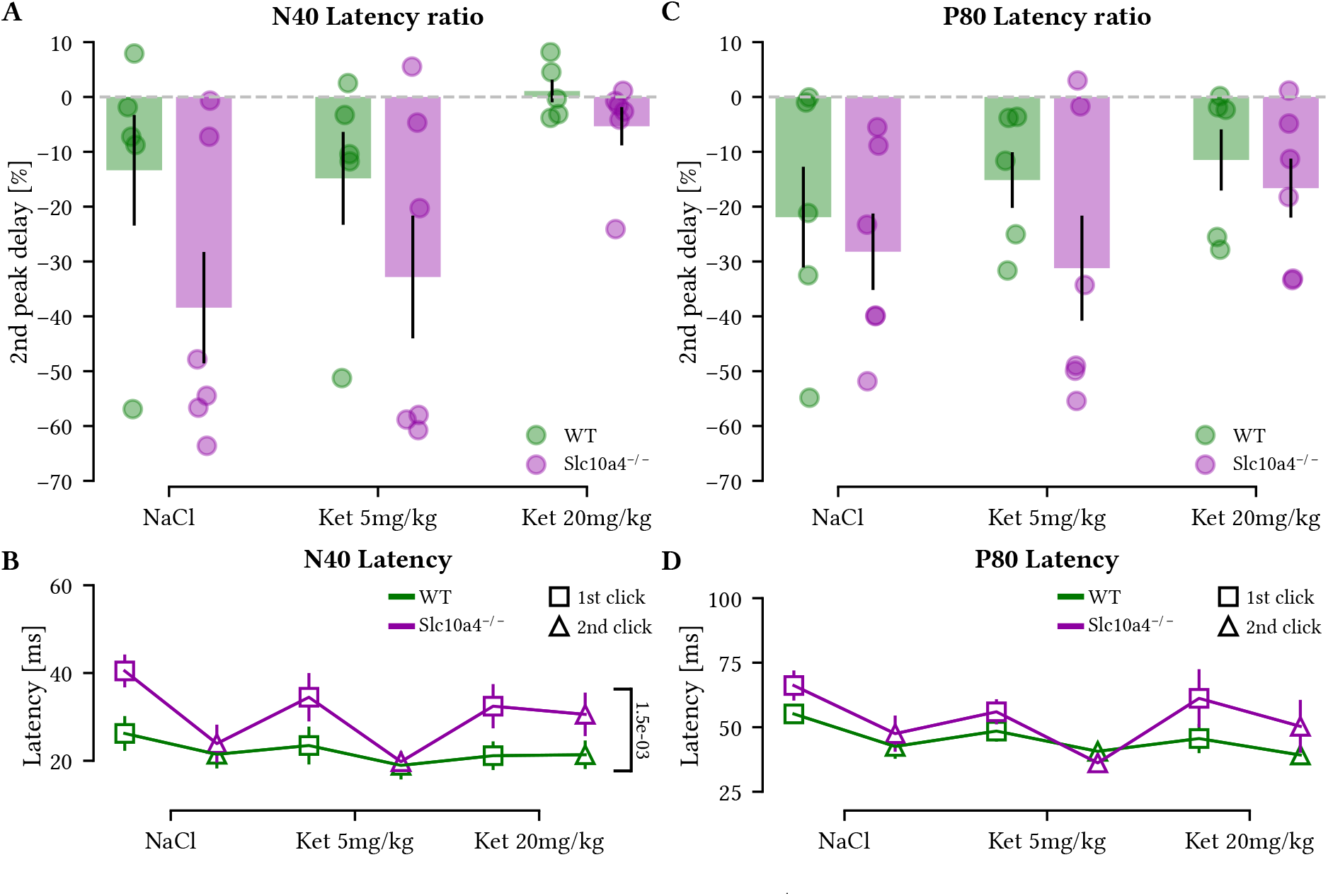
Latency of auditory N40 ERPs is slower in *Slc10a4*^-/-^ mice compared to wildtype littermates. A) Percentage of average second N40 peak delay in relation to the first click for WT (green) and *Slc10a4* ^-/-^ (blue) at each treatment showing no difference between groups or treatments. B) Group average of N40 latency for WT (green) and *Slc10a4* ^-/-^ mice (purple) for each click at each pharmacological treatment showing consistently increased in N40 latency for *Slc10a4* ^-/-^ animals (p = 1.5e-03, Kruskal-Wallis test). C) Percentage of the second P80 peak delay for WT (green) and *Slc10a4* ^-/-^ mice (purple) at each treatment. D) Group average of P80 latency for WT (green) and *Slc10a4* ^-/-^ mice (purple) for each click at each pharmacological treatment, dashed gray line indicates 0% delay. n = 5 WT and 6 *Slc10a4* ^-/-^ mice.

Despite the lack of difference in baseline locomotion between WT and *Slc10a4* ^-/-^ mice, we still evaluated possible sensorimotor gating deficiency in early stage information processing and used a separated group of mice for testing prepulse inhibition. The pre-pulse stimulus intensity varied between 75 or 85 dB, with duration of 4 ms, while the subsequent startle pulse was presented at 105 dB, with duration of 20 ms. The ISI between pre-pulse and startle pulse were 30 or 100 ms (Figure 7A). When the startle amplitude response was delivered alone, no group differences in the performance were observed (Figure 7B). Interestingly, we found interaction effects of genotype with stimulus intensity (ANOVA, F(1,14) = 11.273, p = 5.0e-03) and interstimulus interval (ANOVA, F(1,14) = 10.496, p = 6.0e-03). In agreement with the observed effects, *Slc10a4* ^-/-^ mice displayed no significant differences compared to control littermates using 30 ms of ISI at either intensity (Figure 7C-D), nor using 100 ms of ISI when pre-pulse stimuli was presented at 75 dB (Figure 7E); however, when presented at 85 dB, *Slc10a4* ^-/-^ animals showed significantly decreased startle response compared to WT animals (n = 8 *Slc10a4* ^-/-^ mice and 8 WT mice; Tukey’s HSD eff. size = −1.4; p = 2.4e-02, Figure 7F). These findings suggest a dysfunction in the sensorimotor gating mechanism of *Slc10a4* ^-/-^ animals. Since the decrease in prepulse inhibition was only seen with a longer ISI this indicates that temporal processing is indeed perturbed in *Slc10a4* ^-/-^ mice, compared to WT littermates.

**Figure 7:**
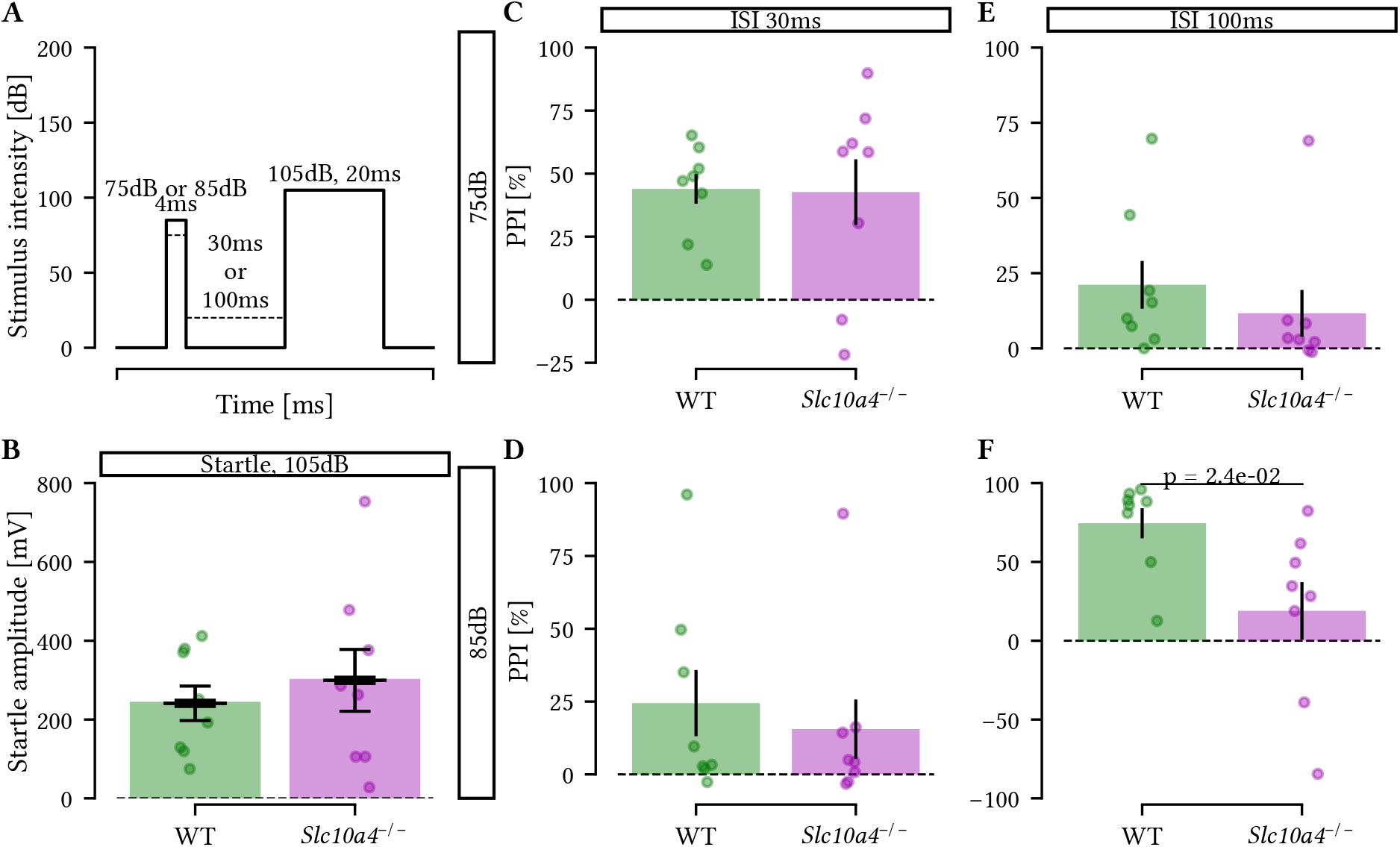
*Slc10a4*^-/-^ mice show less prepulse inhibition with longer interstimulus intervals. A) The protocol used for the PPI testing. Prepulse trials consisted of a single 105 dB pulse preceded by either a 30-ms or a 100-ms inter-stimulus interval (ISI) by a 4-ms non-startling stimulus of 75 or 85 dB. B) The first and last trials of the PPI protocol consisted of single 20 ms 105 dB pulse alone startle stimuli. There was no difference between *Slc10a4* ^-/-^ and WT mice in terms of startle amplitude (mean ± sem). C-F) Four different PPI arrangements tested. The average PPI of startle for the 75 dB pre-pulse, 30-ms inter-stimulus interval (ISI) (C); the 85 dB pre-pulse, 30-ms ISI (D); the 75 dB pre-pulse, 100-ms ISI session (E) and the 85 dB pre-pulse, 100-ms ISI (F) arrangement are shown. Averages and responses from individual mice (n = 8 WT and 8 *Slc10a4* ^-/-^) are shown. Error bars represent SEM.

## Discussion

This study aimed to investigate the effects of a *Slc10a4* gene deletion on auditory sensory gating and sensorimotor gating in mice. Differently from controls, *Slc10a4* ^-/-^ mice showed no amplitude differences between the N40 first and second click in response to paired clicks in saline and ketamine conditions. However, compared to control littermates, *Slc10a4* ^-/-^ mice consistently demonstrated longer N40 latencies and increased P80 amplitudes. Although there were no significant changes in N40 or P80 ratio, sensory motor gating was impaired in *Slc10a4* ^-/-^ animals. These findings suggest that *Slc10a4* ^-/-^ mice exhibit altered auditory processing and altered sensorimotor gating compared to controls, features commonly observed in schizophrenia.

Schizophrenia and psychosis-like states are complex neurological conditions with multiple theories attempting to explain their underlying causes. Two of the most prominent theories are the dopamine theory and the glutamate theory. However, there is a debate in the scientific community as to whether glutamate hypofunction is the primary cause or a secondary effect of dopamine hyperfunction or dysregulation (Laruelle et al, 2005; Lester et al, 2010). Unlike mood disorders where pharmacological treatments targeting specific neurotransmitter systems are available (Mathew et al, 2008), there is no specific pharmacology for treating complex schizophrenia symptoms (Kim et al, 2009). In addition to dopamine, acetylcholine may also be important for the pathophysiology of schizophrenia, since acetylcholinesterase inhibitors have been effective in treating psychosis in Alzheimer’s and Parkinson’s patients (Lester et al, 2010). Therefore, dysfunction in cholinergic neurotransmission may also be associated with schizophrenia.

Our findings are in line with previous research demonstrating that ketamine impairs sensory processing in both animals and humans. Specifically, (Yi et al, 2022) reported that a subanesthetic dose of ketamine decreased the mismatch negativity in mice, a measure of auditory sensory processing. This effect was attributed to the inhibition of GABAergic neurons in the dorsal hippocampus projecting to the medial entorhinal cortex (Yi et al, 2022). Moreover, the study also found increased c-fos staining in the dorsal hippocampus, medial entorhinal and auditory cortices indicating the involvement of these areas in auditory mismatch detection (Yi et al, 2022). Consistent with these findings, we observed that ketamine administration (20 mg/kg) eliminated the difference between the N40 first click and second click amplitude response in wildtype mice, suggesting that this dose affects sensory processing. Notably, the reduction in N100 (equivalent to mice N40) click 1 amplitude (Gjini et al, 2010; Rosburg, 2018) and click 2/click 1 ratio (Gjini et al, 2010) are physiological abnormalities seen in schizophrenia. Instead, *Slc10a4* ^-/-^ mice displayed a lack of a consistent N40 second click amplitude reduction compared to the first click across all treatments, particularly after 5 mg/kg ketamine administration, suggesting that SLC10A4 deficiency may result in less robust sensory processing or filtering in the hippocampal network. However, we did not observe any changes in the auditory sensory gating ratio when comparing WT with *Slc10a4* ^-/-^ mice, indicating that SLC10A4 protein deficiency is compensated for during sensory gating of the N40 component.

The P200 (equivalent to mice P80) component amplitude is thought to reflect attentional allocation and initial conscious awareness of a stimulus, with P200 latency reflecting the temporal aspect of attention (Lijffijt et al, 2009). We observed a trend towards increased, P80 second click amplitude response in the saline condition in *Slc10a4* ^-/-^ mice, possibly indicating excessive sensory processing of this component. Moreover, *Slc10a4* ^-/-^ mice exhibited increased N40 latency and P80 amplitude compared to WT littermates (Figure 8A-D), suggesting impairments in both the first and second click response, respectively.

**Figure 8:**
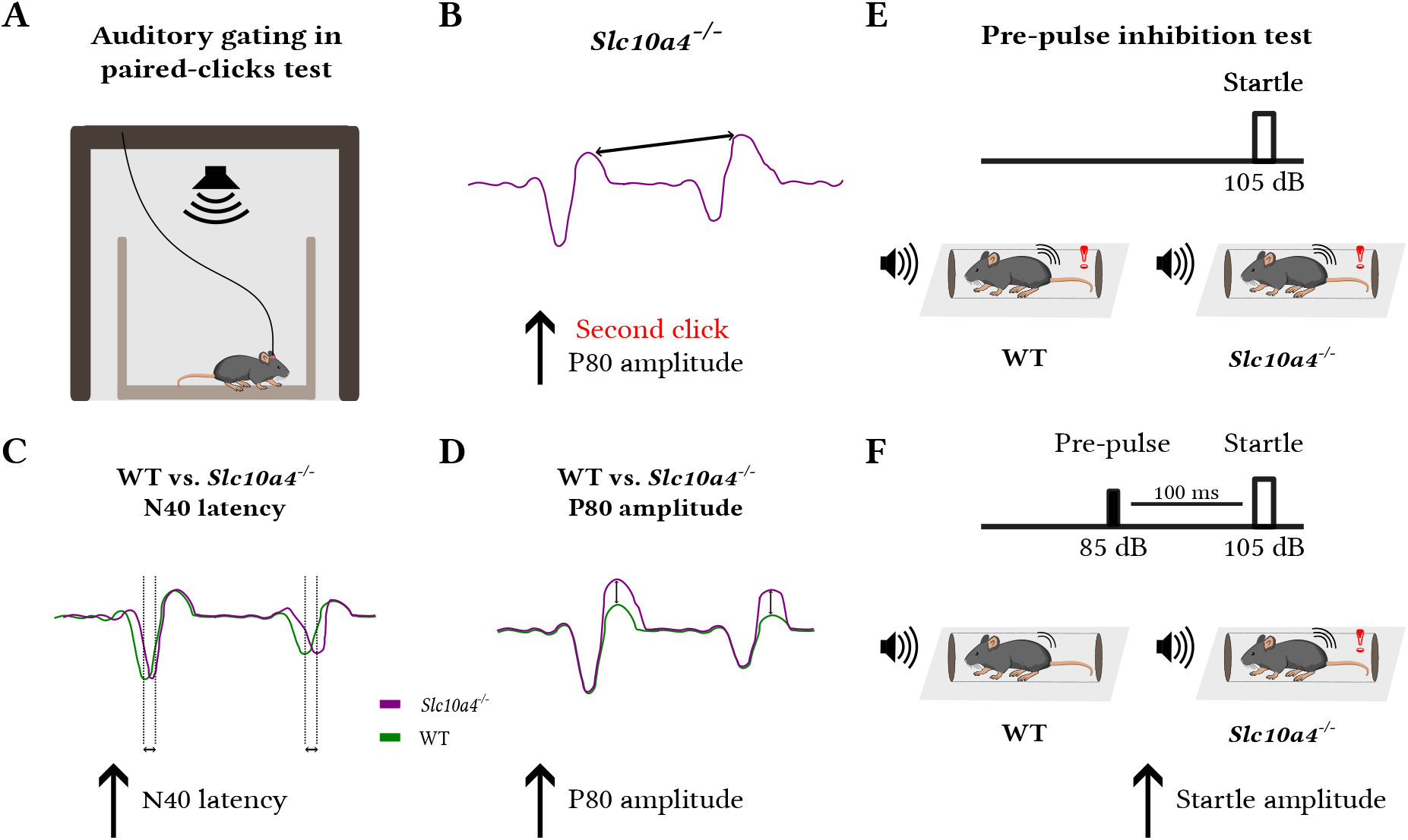
Schematics of the main findings. A) Experimental setup showing an implanted animal during the paired-click test recording. B) P80 second click amplitude tended to be increased compared to the first click response in *Slc10a4* ^-/-^ mice. C) N40 latency is increased in *Slc10a4* ^-/-^ mice compared to control littermates. D) P80 amplitude is increased in *Slc10a4* ^-/-^ mice compared to controls. E) WT and *Slc10a4* ^-/-^ mice showed similar responses in trials with only the startle pulse during PPI test. F) *Slc10a4* ^-/-^ mice present decreased inhibition of startle response in PPI when the interstimulus interval was separated by 100 ms.

As previously reported, *Slc10a4* ^-/-^ mice exhibit altered dopamine and cholinergic signaling with decreased dopamine in the lateral and dorsal striatum and reduced acetylcholine in the brainstem and hippocampus (Melief et al, 2016; Larhammar et al, 2015). Moreover, SLC10A4 deficiency increases susceptibility to cholinergic drugs and epileptic disorders (Zelano et al, 2013). The cholinergic system is known to play a role in the regulation of early aERP components, such as the P20 and N40. Nicotine can improve sensory gating by increasing the amplitude of the first P20 click, an effect that is blocked by DH*β*E (Dihydro-*β*-erythroidine hydrobromide), an antagonist of nicotinic acetylcholine receptor subtypes *α*4/*β*2 and probably *α*2/*β*2 as well (Featherstone et al. 2012). Similarly, nicotine has been shown to improve gating of the N40 component by increasing the first click amplitude, which partly depend on *α*7 receptors, since the antagonist Methyllycaconitine can reduce gating (Rudnick et al, 2010; Featherstone et al, 2012). Moreover, EEG recordings in rats suggest that the N50 (equivalent to the N40 in mice) is sensitive to scopolamine, a non-selective muscarinic antagonist that disrupts sensory gating by increasing the amplitude of the second click (Klinkenberg et al, 2011). These findings suggest that nicotinic acetylcholine receptors regulate the first click, whereas muscarinic receptors modulate the second click of the N40 component.

The sensorimotor gating is also an information processing measurement, and PPI deficits have been repeatedly reported in patients with schizophrenia (Braff et al, 2001; Hedberg et al, 2021; San-Martin et al, 2022). Although some studies show PPI deficits using 30 ms of interstimulus interval (Minassian et al, 2007), our results are similar to Meincke et al (2004) and San-Martin et al (2022), where they observed alterations in schizophrenia patients when the PPI were presented with 100 ms or 120 ms interstimulus, but not in the trials using an ISI of 30 ms (Figure 8E-F). This suggests that *Slc10a4* ^-/-^ animals present increased processing to long-latency repetitive stimulus, which was also observed by the increased second click P80 component, indicating alteration in attenuation of repetitive sensory information.

The monoaminergic and glutamatergic systems have been also implicated in sensory gating deficits. For instance, it has been demonstrated that the dopamine agonist apomorphine can disrupt PPI in rats, which mimics PPI deficits observed in individuals with schizophrenia (Mansbach et al, 1988). Moreover, the specific D2 dopamine receptor antagonist, haloperidol, prevents the effects of apomorphine on prepulse inhibition (Mansbach et al, 1988; Swerdlow, 1994). Similarly, the NMDA receptor antagonist MK-801 disrupts auditory sensory gating. Conversely, selective lesioning of the noradrenergic input with the neurotoxin DSP4 (N-(2-chloroethyl)-N-ethyl-2-bromobenzylamine hydrochloride), as well as less selective depletion of monoamines with reserpine, can block the loss of gating caused by MK-801 (Miller et al, 1992). Taken together, these findings suggest that alterations in cholinergic and monoaminergic transmission in *Slc10a4* ^-/-^ mice may contribute to the observed deficits in sound processing.

In summary, our findings suggest that *Slc10a4* ^-/-^ mice display characteristics present in psychiatric disorders such as schizophrenia due to impaired sensorimotor gating and overprocessing of auditory repeated stimuli (indicated by increased response to the repeated stimulus in P80). Deficient inhibition of P50 (equivalent to the P20 in mice) have been consistently found in schizophrenia patients in previous research (Brockhaus-Dumke et al, 2008; Shen et al, 2020; Freedman et al, 2020), and N100 and P200 amplitude ratio was also shown to be impaired (Gjini et al, 2010; Brockhaus-Dumke et al, 2008; Shen et al, 2020), but according to a meta-analysis involving 1027 schizophrenia patients and 1131 controls, the impairment in ratio of N100 in patients was probably caused by their lower amplitude to the conditioning stimuli (S1) rather than insufficient suppression of the test stimuli (S2), indicating that they weren’t caused by schizophrenia’s sensory gating problems (Rosburg, 2018). Conversely, higher P200 amplitudes have been reported in schizophrenia patients (Ogura et al, 1991). In addition, the PPI phenotype was suggested to be useful for identifying genes related to schizophrenia (Powell et al, 2009). Hence, the *Slc10a4* ^-/-^ animals may serve as a model for investigating how imbalances in brain chemistry contribute to the pathophysiology of psychiatric disorders, given that the altered monoamine and cholinergic transmission could result in auditory and motor sensory processing impairment. A deeper investigation will be needed to clarify this relationship. Of note, the *Slc10a4* gene has been identified as a potential candidate gene in the mapping of genomic loci that implicate synaptic biology in schizophrenia (Trubetskoy et al, 2022). Additionally, a large-scale genome-wide combined analysis of common genetic variants shared among five major psychiatric disorders also identified the Slc10A4 gene as a potential candidate (Xia et al, 2019).

The results of this study suggest that *Slc10a4* ^-/-^ mice exhibit altered auditory processing and sensory motor gating compared to control littermates, which are both features commonly observed in schizophrenia. However, the study alone cannot fully warrant the use of *Slc10a4* ^-/-^ mice in schizophrenia research. Additional research is needed to determine the potential relevance of *Slc10a4* the pathophysiology of schizophrenia and to further investigate the translational potential of this animal model for studying neuropsychiatric disorders.

## Author Contributions

KEL designed the study; KEL, MMH and RNL did experiments; BC and TM analyzed the data; BC and KEL wrote the manuscript with important input from CRC and KK.

## Acknowledgments

The authors would like to thank Dr Sanja Mikulovic for technical advice and electrode tract images; Dr Helton Maia for analysis script improvements; and the communities behind the multiple open-source software packages on which our analysis rely on.

## Data Availability Statement

The datasets generated and/or analyzed in the current study are available on request.

## References

Afshari P, Yao WD, Middleton FA (2017) Reduced slc1a1 expression is associated with neuroinflammation and impaired sensorimotor gating and cognitive performance in mice: Implications for schizophrenia. PLoS One 12(9):e0183,854

Beck K, Hindley G, Borgan F, Ginestet C, McCutcheon R, Brugger S, Driesen N, Ranganathan M, D’Souza DC, Taylor M, Krystal JH, Howes OD (2020) Association of ketamine with psychiatric symptoms and implications for its therapeutic use and for understanding schizophrenia. JAMA Network Open 3(5):e204,693, DOI 10.1001/jamanetworkopen.2020.4693, URL https://doi.org/10.1001/jamanetworkopen.2020.4693

Bickel S, Lipp HP, Umbricht D (2007) Early auditory sensory processing deficits in mouse mutants with reduced nmda receptor function. Neuropsychopharmacology 33(7):1680–1689, DOI 10.1038/sj.npp.1301536, URL http://dx.doi.org/10.1038/sj.npp.1301536

Braff DL, Geyer MA, Light GA, Sprock J, Perry W, Cadenhead KS, Swerdlow NR (2001) Impact of prepulse characteristics on the detection of sensorimotor gating deficits in schizophrenia. Schizophrenia Research 49(1-2):171–178, DOI 10.1016/s0920-9964(00)00139-0, URL http://dx.doi.org/10.1016/s0920-9964(00)00139-0

Brockhaus-Dumke A, Schultze-Lutter F, Mueller R, Tendolkar I, Bechdolf A, Pukrop R, Klosterkoetter J, Ruhrmann S (2008) Sensory gating in schizophrenia: P50 and n100 gating in antipsychotic-free subjects at risk, first-episode, and chronic patients. Biological Psychiatry 64(5):376–384, DOI 10.1016/j.biopsych.2008.02.006, URL https://doi.org/10.1016%2Fj.biopsych.2008.02.006

Burger S, Däring B, Hardt M, Beuerlein K, Gerstberger R, Geyer J (2011) Co-expression studies of the orphan carrier protein slc10a4 and the vesicular carriers vacht and vmat2 in the rat central and peripheral nervous system. Neuroscience 193:109–121, DOI 10.1016/j.neuroscience.2011.06.068, URL https://doi.org/10.1016/j.neuroscience.2011.06.068

van den Buuse M (2009) Modeling the positive symptoms of schizophrenia in genetically modified mice: Pharmacology and methodology aspects. Schizophrenia Bulletin 36(2):246–270, DOI 10.1093/schbul/sbp132, URL https://doi.org/10.1093/schbul/sbp132

Chatterjee M, Ganguly S, Srivastava M, Palit G (2011) Effect of ‘chronic’ versus ‘acute’ ketamine administration and its ‘withdrawal’ effect on behavioural alterations in mice: Implications for experimental psychosis. Behavioural Brain Research 216(1):247–254, DOI 10.1016/j.bbr.2010.08.001, URL https://doi.org/10.1016/j.bbr.2010.08.001

de Chaumont F, Dallongeville S, Chenouard N, Hervé N, Pop S, Provoost T, Meas-Yedid V, Pankajakshan P, Lecomte T, Montagner YL, Lagache T, Dufour A, Olivo-Marin JC (2012) Icy: an open bioimage informatics platform for extended reproducible research. Nature Methods 9(7):690–696, DOI 10.1038/nmeth.2075, URL https://doi.org/10.1038/nmeth.2075

Ciralli B, Malfatti T, Lima TZ, Silva SRB, Cederroth CR, Leao KE (2022) Alterations of auditory sensory gating in mice with noise-induced tinnitus treated with nicotine and cannabis extract. bioRxiv DOI 10.1101/2022.06.18.496668, URL https://doi.org/10.1101/2022.06.18.496668

Connolly PM, Maxwell C, Liang Y, Kahn JB, Kanes SJ, Abel T, Gur RE, Turetsky BI, Siegel SJ (2004) The effects of ketamine vary among inbred mouse strains and mimic schizophrenia for the p80, but not p20 or n40 auditory erp components. Neurochemical Research 29(6):1179–1188, DOI 10.1023/b:nere.0000023605.68408.fb, URL http://dx.doi.org/10.1023/b:nere.0000023605.68408.fb

Creese I, Burt DR, Snyder SH (1976) Dopamine receptor binding predicts clinical and pharmacological potencies of antischizophrenic drugs. Science 192(4238):481–483

Ehrlichman R, Gandal M, Maxwell C, Lazarewicz M, Finkel L, Contreras D, Turetsky B, Siegel S (2009) N-methyl-d-aspartic acid receptor antagonist–induced frequency oscillations in mice recreate pattern of electrophysiological deficits in schizophrenia. Neuroscience 158(2):705–712, DOI 10.1016/j.neuroscience.2008.10.031, URL https://doi.org/10.1016/j.neuroscience.2008.10.031

Featherstone RE, Phillips JM, Thieu T, Ehrlichman RS, Halene TB, Leiser SC, Christian E, Johnson E, Lerman C, Siegel SJ (2012) Nicotine receptor subtype-specific effects on auditory evoked oscillations and potentials. PLoS ONE 7(7):e39,775, DOI 10.1371/journal.pone.0039775, URL https://doi.org/10.1371/journal.pone.0039775

Freedman R, Olsen-Dufour AM, Olincy A (2020) P50 inhibitory sensory gating in schizophrenia: analysis of recent studies. Schizophrenia Research 218:93–98, DOI 10.1016/j.schres.2020.02.003, URL https://doi.org/10.1016/j.schres.2020.02.003

Geier M (2021) svgutils. https://pypi.org/project/svgutils/, software

Geyer J, Fernandes C, Döring B, Burger S, Godoy J, Rafalzik S, Hübschle T, Gerstberger R, Petzinger E (2008) Cloning and molecular characterization of the orphan carrier protein slc10a4: Expression in cholinergic neurons of the rat central nervous system. Neuroscience 152(4):990–1005, DOI 10.1016/j.neuroscience.2008.01.049, URL https://doi.org/10.1016/j.neuroscience.2008.01.049

Gjini K, Arfken C, Boutros NN (2010) Relationships between sensory “gating out” and sensory “gating in” of auditory evoked potentials in schizophrenia: A pilot study. Schizophrenia Research 121(1-3):139–145, DOI 10.1016/j.schres.2010.04.020, URL https://doi.org/10.1016%2Fj.schres.2010.04.020

Harris CR, Millman KJ, van der Walt SJ, Gommers R, Virtanen P, Cournapeau D, Wieser E, Taylor J, Berg S, Smith NJ, Kern R, Picus M, Hoyer S, van Kerkwijk MH, Brett M, Haldane A, Fernández del Río J, Wiebe M, Peterson P, Gérard-Marchant P, Sheppard K, Reddy T, Weckesser W, Abbasi H, Gohlke C, Oliphant TE (2020) Array programming with NumPy. Nature 585:357–362, DOI 10.1038/s41586-020-2649-2

Hedberg M, Imbeault S, Erhardt S, Schwieler L (2021) Disrupted sensorimotor gating in first-episode psychosis patients is not affected by short-term antipsychotic treatment. Schizophrenia Research 228:118–123, DOI 10.1016/j.schres.2020.12.009, URL https://doi.org/10.1016%2Fj.schres.2020.12.009

Hunter JD (2007) Matplotlib: A 2d Graphics Environment. Computing in Science & Engineering 9(3):90–95, DOI 10.1109/MCSE.2007.55, URL http://ieeexplore.ieee.org/document/4160265/

Inkscape Project (2022) Inkscape. URL https://inkscape.org, software

Javitt DC, Sweet RA (2015) Auditory dysfunction in schizophrenia: integrating clinical and basic features. Nature Reviews Neuroscience 16(9):535–550, DOI 10.1038/nrn4002, URL https://doi.org/10.1038/nrn4002

Kim DH, Maneen MJ, Stahl SM (2009) Building a better antipsychotic: Receptor targets for the treatment of multiple symptom dimensions of schizophrenia. Neurotherapeutics 6(1):78–85, DOI 10.1016/j.nurt.2008.10.020, URL http://dx.doi.org/10.1016/j.nurt.2008.10.020

Klinkenberg I, Sambeth A, Blokland A (2011) Acetylcholine and attention. Behavioural brain research 221(2):430–442

Kristensen AS, Andersen J, Jørgensen TN, Sørensen L, Eriksen J, Loland CJ, Strømgaard K, Gether U (2011) Slc6 neurotransmitter transporters: Structure, function, and regulation. Pharmacological Reviews 63(3):585–640, DOI 10.1124/pr.108.000869, URL https://doi.org/10.1124/pr.108.000869

Krystal JH, Karper LP, Seibyl JP, Freeman GK, Delaney R, Bremner JD, Heninger GR, Bowers MB, Charney DS (1994) Subanesthetic effects of the noncompetitive nmda antagonist, ketamine, in humans: psychotomimetic, perceptual, cognitive, and neuroendocrine responses. Archives of general psychiatry 51(3):199–214

Larhammar M, Patra K, Blunder M, Emilsson L, Peuckert C, Arvidsson E, Rönnlund D, Preobraschenski J, Birgner C, Limbach C, Widengren J, Blom H, Jahn R, Wallén-Mackenzie Å, Kullander K (2015) SLC10a4 is a vesicular amine-associated transporter modulating dopamine homeostasis. Biological Psychiatry 77(6):526–536, DOI 10.1016/j.biopsych.2014.07.017, URL https://doi.org/10.1016/j.biopsych.2014.07.017

Laruelle M, Frankle WG, Narendran R, Kegeles LS, Abi-Dargham A (2005) Mechanism of action of antipsychotic drugs: from dopamine d2 receptor antagonism to glutamate nmda facilitation. Clinical therapeutics 27:S16–S24

Lester DB, Rogers TD, Blaha CD (2010) Acetylcholine-dopamine interactions in the pathophysiology and treatment of cns disorders. CNS Neuroscience & Therapeutics 16(3):137–162, DOI 10.1111/j.1755-5949.2010.00142.x, URL https://doi.org/10.1111/j.1755-5949.2010.00142.x

Lijffijt M, Lane SD, Meier SL, Boutros NN, Burroughs S, Steinberg JL, Moeller FG, Swann AC (2009) P50, n100, and p200 sensory gating: Relationships with behavioral inhibition, attention, and working memory. Psychophysiology 46(5):1059–1068, DOI 10.1111/j.1469-8986.2009.00845.x, URL https://doi.org/10.1111/j.1469-8986.2009.00845.x

Lin L, Yee SW, Kim RB, Giacomini KM (2015) Slc transporters as therapeutic targets: emerging opportunities. Nature Reviews Drug Discovery 14(8):543–560, DOI 10.1038/nrd4626, URL https://doi.org/10.1038/nrd4626

Malfatti T (2023) Sciscripts. DOI 10.5281/ZENODO.4045872, URL https://zenodo.org/record/4045872, software

Malfatti T, Ciralli B (2023) Sensorygatingonslc10a4. URL https://gitlab.com/malfatti/SensoryGatingOnSLC10A4, software repository

Mansbach RS, Geyer MA, Braff DL (1988) Dopaminergic stimulation disrupts sensorimotor gating in the rat. Psychopharmacology 94(4):507–514

Mathew SJ, Manji HK, Charney DS (2008) Novel drugs and therapeutic targets for severe mood disorders. Neuropsychopharmacology 33(9):2080–2092, DOI 10.1038/sj.npp.1301652, URL http://dx.doi.org/10.1038/sj.npp.1301652

Meincke U, MÃ¶rth D, Voß T, Thelen B, Geyer MA, Gouzoulis–Mayfrank E (2004) Prepulse inhibition of the acoustically evoked startle reflex in patients with an acute schizophrenic psychosis—a longitudinal study. European Archives of Psychiatry and Clinical Neuroscience 254(6):415–421, DOI 10.1007/s00406-004-0523-0, URL https://doi.org/10.1007%2Fs00406-004-0523-0

Melief E, Gibbs J, Li X, Morgan R, Keene C, Montine T, Palmiter R, Darvas M (2016) Characterization of cognitive impairments and neurotransmitter changes in a novel transgenic mouse lacking slc10a4. Neuroscience 324:399–406, DOI 10.1016/j.neuroscience.2016.03.037, URL https://doi.org/10.1016/j.neuroscience.2016.03.037

Miller CL, Bickford PC, Luntz-Leybman V, Adler L, Gerhardt G, Freedman R (1992) Phencyclidine and auditory sensory gating in the hippocampus of the rat. Neuropharmacology 31(10):1041–1048

Minassian A, Feifel D, Perry W (2007) The relationship between sensorimotor gating and clinical improvement in acutely ill schizophrenia patients. Schizophrenia Research 89(1-3):225–231, DOI 10.1016/j.schres.2006.08.006, URL https://doi.org/10.1016%2Fj.schres.2006.08.006

Ogura C, Nageishi Y, Matsubayashi M, Omura F, Kishimoto A, Shimokochi M (1991) Abnormalities in event-related potentials, n100, p200, p300 and slow wave in schizophrenia. Psychiatry and Clinical Neurosciences 45(1):57–65, DOI 10.1111/j.1440-1819.1991.tb00506.x, URL http://dx.doi.org/10.1111/j.1440-1819.1991.tb00506.x

Patra K, Lyons DJ, Bauer P, Hilscher MM, Sharma S, Leão RN, Kullander K (2014) A role for solute carrier family 10 member 4, or vesicular aminergic-associated transporter, in structural remodelling and transmitter release at the mouse neuromuscular junction. European Journal of Neuroscience 41(3):316–327, DOI 10.1111/ejn.12790, URL https://doi.org/10.1111/ejn.12790

Pizzagalli MD, Bensimon A, Superti-Furga G (2020) A guide to plasma membrane solute carrier proteins. The FEBS Journal 288(9):2784–2835, DOI 10.1111/febs.15531, URL https://doi.org/10.1111/febs.15531

Powell SB, Zhou X, Geyer MA (2009) Prepulse inhibition and genetic mouse models of schizophrenia. Behavioural Brain Research 204(2):282–294, DOI 10.1016/j.bbr.2009.04.021, URL https://doi.org/10.1016%2Fj.bbr.2009.04.021

Rosburg T (2018) Auditory n100 gating in patients with schizophrenia: A systematic meta-analysis. Clinical Neurophysiology 129(10):2099–2111, DOI 10.1016/j.clinph.2018.07.012, URL https://doi.org/10.1016%2Fj.clinph.2018.07.012

Rudnick ND, Strasser AA, Phillips JM, Jepson C, Patterson F, Frey JM, Turetsky BI, Lerman C, Siegel SJ (2010) Mouse model predicts effects of smoking and varenicline on event-related potentials in humans. Nicotine & Tobacco Research 12(6):589–597, DOI 10.1093/ntr/ntq049, URL https://doi.org/10.1093/ntr/ntq049

San-Martin R, Zimiani M, de Avila M, Shuhama R, Del-Ben C, Menezes P, Fraga F, Salum C (2022) Early schizophrenia and bipolar disorder patients display reduced neural prepulse inhibition. Brain Sciences 12(1):93, DOI 10.3390/brainsci12010093, URL http://dx.doi.org/10.3390/brainsci12010093

Scheffer-Teixeira R, Tort ABL (2017) Unveiling fast field oscillations through comodulation. eneuro 4(4):ENEURO.0079–17.2017, DOI 10.1523/eneuro.0079-17.2017, URL https://doi.org/10.1523/eneuro.0079-17.2017

Schmidt S, Moncada M, Burger S, Geyer J (2015) Expression, sorting and transport studies for the orphan carrier slc10a4 in neuronal and non-neuronal cell lines and in xenopus laevis oocytes. BMC Neuroscience 16(1), DOI 10.1186/s12868-015-0174-2, URL https://doi.org/10.1186/s12868-015-0174-2

Seeman P, Lee T (1975) Antipsychotic drugs: direct correlation between clinical potency and presynaptic action on dopamine neurons. Science 188(4194):1217–1219

Shen CL, Chou TL, Lai WS, Hsieh MH, Liu CC, Liu CM, Hwu HG (2020) P50, n100, and p200 auditory sensory gating deficits in schizophrenia patients. Frontiers in Psychiatry 11, DOI 10.3389/fpsyt.2020.00868, URL https://doi.org/10.3389/fpsyt.2020.00868

Siegel S, Maxwell C, Majumdar S, Trief D, Lerman C, Gur R, Kanes S, Liang Y (2005) Monoamine reuptake inhibition and nicotine receptor antagonism reduce amplitude and gating of auditory evoked potentials. Neuroscience 133(3):729–738, DOI 10.1016/j.neuroscience.2005.03.027, URL https://doi.org/10.1016/j.neuroscience.2005.03.027

Splinter PL, Lazaridis KN, Dawson PA, LaRusso NF (2006) Cloning and expression of slc10a4, a putative organic anion transport protein. World journal of gastroenterology: WJG 12(42):6797

Swerdlow NR (1994) Assessing the validity of an animal model of deficient sensorimotor gating in schizophrenic patients. Archives of General Psychiatry 51(2):139, DOI 10.1001/archpsyc.1994.03950020063007, URL http://dx.doi.org/10.1001/archpsyc.1994.03950020063007

Trubetskoy V, Pardiñas A, Qi T (2022) Psychencode; psychosis endophenotypes international consortium; syngo consortium; schizophrenia working group of the psychiatric genomics consortium. mapping genomic loci implicates genes and synaptic biology in schizophrenia. Nature 604(7906):502–508, DOI 10.1038/s41586-022-04434-5

Virtanen P, Gommers R, Oliphant TE, Haberland M, Reddy T, Cournapeau D, Burovski E, Peterson P, Weckesser W, Bright J, van der Walt SJ, Brett M, Wilson J, Millman KJ, Mayorov N, Nelson ARJ, Jones E, Kern R, Larson E, Carey CJ, Polat I, Feng Y, Moore EW, VanderPlas J, Laxalde D, Perktold J, Cimrman R, Henriksen I, Quintero EA, Harris CR, Archibald AM, Ribeiro AH, Pedregosa F, van Mulbregt P, SciPy 10 Contributors (2020) SciPy 1.0: Fundamental Algorithms for Scientific Computing in Python. Nature Methods 17:261–272, DOI 10.1038/s41592-019-0686-2

Winship IR, Dursun SM, Baker GB, Balista PA, Kandratavicius L, de Oliveira JPM, Hallak J, Howland JG (2018) An overview of animal models related to schizophrenia. The Canadian Journal of Psychiatry 64(1):5–17, DOI 10.1177/0706743718773728, URL https://doi.org/10.1177/0706743718773728

Xia L, Xia K, Weinberger D, Zhang F (2019) Common genetic variants shared among five major psychiatric disorders: a large-scale genome-wide combined analysis. Global Clinical and Translational Research pp 21–30, DOI 10.36316/gcatr.01.0003, URL http://dx.doi.org/10.36316/gcatr.01.0003

Yi GL, Zhu MZ, Cui HC, Yuan XR, Liu P, Tang J, Li YQ, Zhu XH (2022) A hippocampus dependent neural circuit loop underlying the generation of auditory mismatch negativity. Neuropharmacology 206:108,947, DOI 10.1016/j.neuropharm.2022.108947, URL https://doi.org/10.1016/j.neuropharm.2022.108947

Zelano J, Mikulovic S, Patra K, Kühnemund M, Larhammar M, Emilsson L, Leao R, Kullander K (2013) The synaptic protein encoded by the gene slc10a4 suppresses epileptiform activity and regulates sensitivity to cholinergic chemoconvulsants. Experimental Neurology 239:73–81, DOI 10.1016/j.expneurol.2012.09.006, URL https://doi.org/10.1016/j.expneurol.2012.09.006

